# Maternal Care Remains Stable in a Predator Odor Model of Early Life Adversity

**DOI:** 10.64898/2026.11.13.766531

**Authors:** Victoria R. Riesgo, Margie C. Stricklin, Hannah E. Carney, Judy E. Nyakoa, Cecelia J. Crandall, Debra A. Bangasser, Amelia Cuarenta

## Abstract

Early life adversity (ELA) creates lasting impacts on the brain and behavior that contribute to vulnerability for psychiatric disorders. Most rodent models of ELA involve manipulating maternal care itself (e.g., maternal separation) or involve changes to the environment that alter maternal care (e.g., low resources). Here, we used neonatal predator odor exposure (POE) as a threat model of ELA in rats where pups themselves are exposed to stressful odors in the absence of the dam. Our previous research shows that POE results in long-term behavioral effects, including reduced social play, but we have yet to determine whether POE also alters maternal care in ways that could contribute to these outcomes. To investigate this, litters were removed from their home cage and exposed to either POE or control conditions from postnatal days 1-3. Immediately following this exposure, pups were returned to their home cage and placed on the side opposite of the nest. Retrieval of pups to the nest and maternal care behaviors were recorded for 45 minutes. On nearly all measures there were no differences in maternal behaviors between POE and control dams. Some behaviors changed across days similarly in POE and control groups: time spent licking and grooming pups increased, and time spent nest building and rearing decreased across days. The only significant difference between conditions was observed on day 1, with control dams spending more time carrying pups during retrieval but this decreased to POE levels on days 2 and 3. Ultimately, maternal care behavior remains largely unchanged by POE. These findings provide evidence that long-lasting effects of POE on offspring are due to POE itself, and not due to changes in maternal care behavior.

**Highlights:**

- Predator Odor Exposure (POE) induces early life adversity.
- Whether POE affects maternal care was previously unknown.
- Maternal care behaviors remained largely stable across POE and control conditions.
- Impacts of POE on later outcomes are not mediated by changes to maternal care.

## Introduction

Early life adversity (ELA) occurs during critical developmental windows where the brain is changing rapidly and sensitive to the influence of experience (Hensch, 2004). ELA has both acute and long-lasting impacts on the brain and behavior. ELA, defined as one or more adverse childhood events, is a common occurrence reported by as many as 76.1% of U.S. high school students (Swedo, 2024). Importantly, ELA is associated with higher vulnerability across the lifespan for psychiatric disorders like anxiety, depression, and substance use disorder (Anda et al., 2006; Green et al., 2010; LeMoult et al., 2020).

Many rodent models of ELA impact both dams and their litters and have been shown to alter maternal care behaviors. Maternal separation, where the dam is manipulated and removed from the litter, produces altered patterns of maternal care including low frequency of care immediately following the separation (Orso et al., 2018). Limited bedding and nesting, a resource scarcity model of ELA, manipulates the environment affecting both the dam and the pups. Limited bedding and nesting can alter time spent nursing and licking and grooming pups and consistently causes maternal care to become fragmented and unpredictable (Baram et al., 2012; Eck et al., 2020; Goodwill et al., 2019; Ivy et al., 2008; Walker et al., 2017). Fragmented maternal care has lasting impacts, including delayed hippocampal impairment in adulthood (Brunson et al., 2005). Altered and unpredictable maternal care patterns, as measured by high behavioral entropy rates, is associated with anhedonia during adolescence (Molet et al., 2016). However, maternal care can also mitigate some effects of ELA, as higher levels of maternal licking and grooming can lessen hypothalamic-pituitary-adrenal axis responses to stress (Liu et al., 1997). Understanding how ELA models influence maternal care can help distinguish the contributions of maternal care and ELA exposure to risk or resilience.

Our lab uses a predator odor exposure (POE) model of ELA, which is a naturalistic and ethologically relevant stressor for our rat model (Kigar et al., 2024). A unique feature of this model is that pups experience perceived predation threat without direct exposure of the dams to the odor. In adult rats, POE produces twice the medial amygdala activation as a foot shock stressor (Baisley et al., 2011), and repeated POE produces a neural and behavioral phenotype similar to that of post-traumatic stress disorder (Rajbhandari et al., 2015), demonstrating POE’s efficacy as a threat model of ELA.

Neonatal POE has both acute and persistent effects on the developing brain. On postnatal day 3, neonatal POE increases the signaling of serotonin in the male and female amygdala (Kigar et al., 2017). At the same age, neonatal POE decreases expression levels of mu- and delta-opioid receptor mRNA in the female nucleus accumbens (Chang et al., 2019), suggesting that some of the mechanistic effects of POE are sex-dependent. The impacts of neonatal POE also extend throughout development, as neonatal POE has been found to decrease both juvenile social play (Cuarenta et al., 2021) and adult social reward motivation (Stricklin et al., 2026).

Neonatal POE also produces both short- and long-term impacts on exposed offspring, but whether dams are able to detect if their pups received POE remains unknown. Although dams do not directly experience POE, they may detect changes or residual odors in exposed pups upon their return to the home cage and alter their maternal care behaviors accordingly. Thus, the effects of POE on the brain and behavior later in life could result from POE itself, subsequent changes in the maternal care, or a combination of these factors. We addressed this research gap by analyzing maternal care behaviors in dams immediately following the return of their pups from POE.

We hypothesized that the dams of POE litters would spend more time engaging in pup-directed behaviors than those of control litters. Although it is unknown whether dams can detect residual predator odor on the pups following POE, we predict that if they can detect the odor or other pup distress cues, they would alter their maternal behavior response. Specifically, dams may spend more time on pup directed behaviors including sniffing and licking and grooming pups in response to the unfamiliar odor. Alternatively, dams may find the predator odor aversive and spend less time engaging in pup-directed behaviors. Thus, changes in maternal care in either direction would be consistent with dams detecting and responding to residual predator odor on their pups. This study advances our characterization of the neonatal POE model by determining whether neonatal POE alters maternal care, as changes in maternal care could contribute to the developmental impacts of early-life threat.

## Methods

### Subjects and predator odor exposure (POE)

Georgia State University Institutional Animal Care and Use Committee approved all animal care and experiments in this project. Sprague Dawley rats were received from Charles River Laboratories (Wilmington, MA) and bred in-house. All POE and control dams were primiparous. Litters were culled to ten pups with a 6F/4M or a 6M/4F composition on postnatal day (PND) 1. One litter contained only nine offspring (5M/4F) due to the dam only giving birth to nine pups. From PND1-3, neonatal rat pups were briefly separated from their home cage and moved into a separate room where they were placed on an elevated, perforated platform inside of an odor exposure chamber. These chambers and platforms were 3D printed on a Bambu X1C (print file <u>linked here</u>). Underneath the platform was a mixture of Bed-o’Cobs bedding (The Andersons) and predator urine. Pups were not directly in contact with the urine-infused bedding. On PND1, the pups were exposed to bobcat urine. On PND2, the pups were exposed to an unfamiliar conspecific male’s urine. On PND3, the pups were exposed to ferret urine. A different predator odor was used each day to avoid habituation. Control litters were handled in the same manner but exposed to clean bedding. The rat pups were monitored for the 5-minute duration of exposure before pups were returned to their home cage. Pups were returned to their home cage in < 15 minutes. Rats were housed in a reverse light/dark cycle with behavioral tasks taking place during the dark cycle between 11:00am and 2:00pm. All rats had *ad libitum* access to water and standard laboratory chow (5001, LabDiet) throughout the duration of the experiment. All rats were housed in standard shoebox cages (20×20×40 cm) with vent filter tops on non-ventilated racks.

### Maternal behavior recordings

Following POE, pups were returned to their home cage, and maternal behavior was recorded for 45 minutes to observe if neonatal POE impacts maternal care behavior. Pups were placed on the opposite side of their home cage from the nesting material to observe pup retrieval behaviors following POE. Cameras were placed over the cages to record from an overhead angle. Cages and dates were labeled in each video with an index card for future scoring. Once the recording started, experimenters exited the room for the 45 minutes to not disrupt maternal behavior. Once the recording period finished, the cage was returned to the animal colony room, and the dams and pups were left undisturbed until the following day of POE.

### Scoring maternal behavior

The 45-minute videos were uploaded to Noldus 15 Observer software and analyzed for both pup retrieval and maternal care behaviors. Two independent researchers blind to condition scored the videos for latency to retrieve all pups to the nest, time spent carrying pups during retrieval, duration of time spent on nest, time spent off nest, pup-directed behaviors, and non-pup directed behaviors. Inter-rater reliability analysis was performed using Noldus Observer’s reliability analysis function. For each video scored, the two researchers were compared for their similarity on both duration and sequence of behaviors, with a comparison cut-off of 1 second and a concordance rate requirement of 80% or higher.

Latency to retrieve all pups to the nest was recorded when the last pup was brought to the nest by the dam. During this time, time spent carrying pups to retrieval was also recorded. ‘On nest’ was defined as beginning when the majority of the dam’s head or backside was over the nest and ending when she moved the majority of her body off the nest and was not interacting with pups. Pup-directed behaviors included archback nursing, passive nursing, blanket nursing, licking and grooming, sniffing, pup carrying, and nest building. Archback nursing was defined as the dam crouched above the pups with fore- and hind-limbs rigid and splayed to the side, with an arched back supporting her own weight while nursing, and some or all pups were visibly attached to dam’s nipples. Passive nursing was defined as the dam lying on her side and not supporting herself with her limbs with one or more pups attached to her nipples. Blanket nursing was defined as the dam being over the litter but without an arched back and no obvious extension/splaying of legs (Myers et al., 1989). Licking and grooming was defined as the dam contacting the pup’s body or anogenital region with her nose and mouth. If the pup dam interaction was occluded, licking and grooming was scored when her head was pointed down and moving in a rhythmic motion consistent with licking and grooming (Moore et al., 1997). Sniffing was defined as the dam moving her nose with rapid and rhythmic respiration proximal to pups. Pup carrying was defined as the dam picking up and carrying a pup with her mouth. Nest building was defined as the dam moving, rearranging, or in any way manipulating the nest.

Non-pup directed behaviors included self-grooming, and rearing. Self-grooming was defined as the dam stroking her body or face with her hands in repetitive elliptical, unilateral, or bilateral strokes or licking her body. Rearing was defined as an instance of the dam lifting two paws off the floor of the cage to lean on the cage wall, which was often accompanied by sniffing or chewing on the cage lid.

Behavioral entropy scores, which are a measurement of behavior fragmentation, were calculated using RStudio based on the start and stop times for each behavior across the 45-minute video. Behavioral entropy provides a quantitative measurement of diversity or unpredictability in the behaviors of the dam, where a higher behavioral entropy score is indicative of more erratic and fragmented behaviors. A total of 12 litters with their dams, 6 POE and 6 control, were observed. Day 3 observations for one control dam were excluded from analyses due to issues with video recording that rendered the video unobservable.

### Statistical analysis

Sample sizes are included in figure legends and reflect the number of litters analyzed. Two-way ANOVAs followed by Tukey post hoc comparisons were used to assess effects of condition on behavior across days. Welch’s unpaired t-tests were used to assess effects of condition on total behavior collapsed across days and can be found in supplementary data. Maternal care entropy scores were calculated using RStudio. All analyses were performed in GraphPad Prism version 11.0.0. Significance thresholds were set to p<0.05.

### Ethical note

The care and use of the animal subjects in this experiment were reviewed and approved by Georgia State University’s Institutional Animal Care and Use Committee (A23020). The purpose of POE is to induce ELA via odor stimuli from natural rodent predators, resulting in anticipated distress that would be antithetical to alleviate. This distress may look like increased mobility from pups and long-term behavioral changes. The time that pups were away from the dam and the home cage was recorded each day of POE and kept under fifteen minutes to reduce any additional stress of maternal separation. Subjects for this experiment were bred from rats obtained from Charles River Laboratories (Wilmington, MA). Six POE litters and five control litters with ten pups each were observed in their home cage. Pups from this experiment were utilized in later experiments evaluating the long-term impacts of POE on behavior.

## Results

### Pup retrieval

There was no significant main effect of condition alone on time spent carrying pups during retrieval (*F* _1, 9_ = 2.088, *p* = 0.1823). There were no significant main effects of day (*F*_1.044, 9.397_ = 0.578, *p* = 0.47), condition (*F*_1, 9_ = 1.277, *p* = 0.29), or a day condition interaction (*F*_1.044, 9.397_ = 0.737, *p* = 0.42) for latency to retrieve pups (Fig. 1A). A two-way ANOVA revealed there was a significant effect of day (*F*_1.153, 10.38_ = 19.74, *p* = 0.0009) and a day ⨯ condition interaction (*F* _1.153, 10.38_ = 19.29, *p* = 0.0009) for the time spent carrying pups during retrieval, with Tukey post hoc comparisons revealing that on day 1, control dams spent more time carrying to retrieve than POE *(p* = 0.0002), and control dams spent significantly more time carrying pups to retrieval on day 1 than days 2 (*p* = 0.0002) and 3 (*p* = 0.0001) (Fig. 1B).

**Figure 1.**
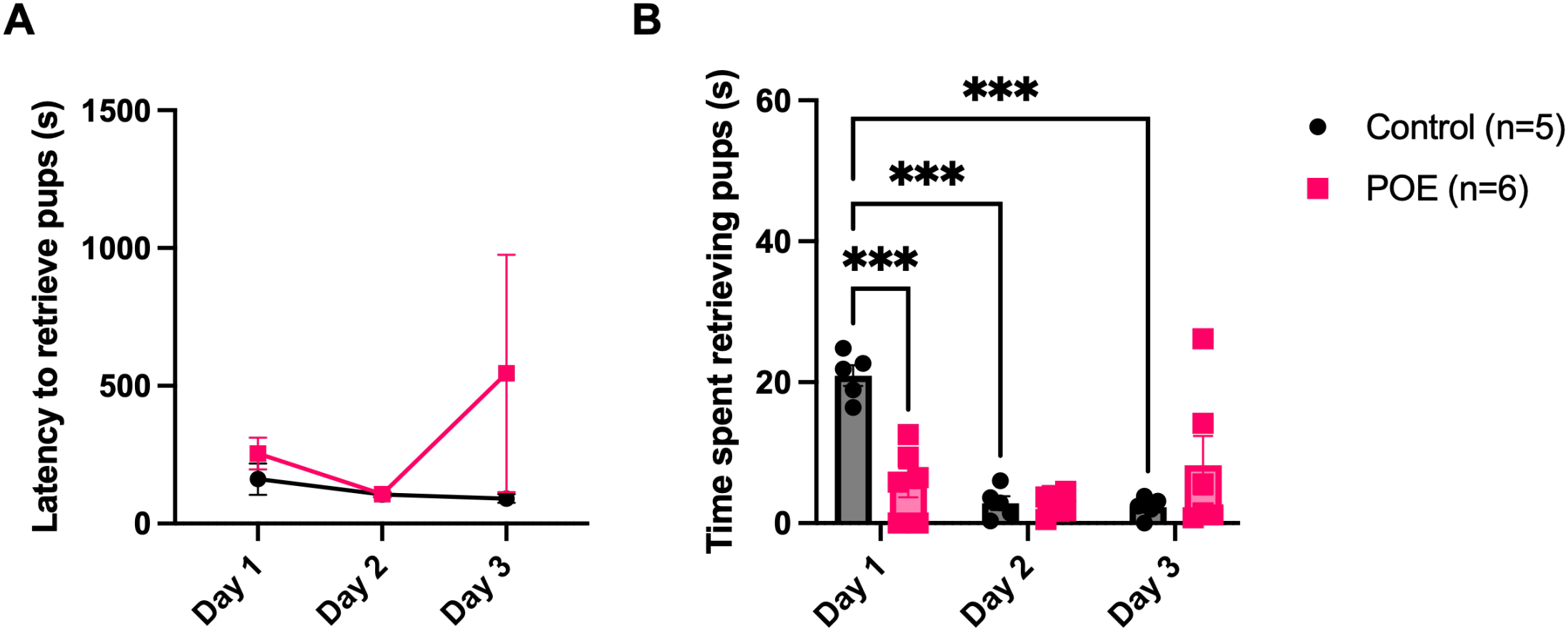
Pup retrieval. A) Latency to retrieve all pups across days. B) Time spent carrying pups to retrieval across days. ***p<.001. Data are represented as means ± SEMS. POE, Predator Odor Exposure.

### Time spent on and off nest

There were no significant main effects of day (*F*_1.571, 14.14_ = 1.767, *p* = 0.78), condition (*F*_1, 9_ = 0.9478, *p* = 0.36), or a day ⨯ condition interaction (*F*_1.571, 14.14_ = 0.1181, *p* = 0.84) for amount of time spent on nest (Fig. 2A). There were no significant main effects of day (*F*_1.574, 14.17_ = 1.899, *p* = 0.19), condition (*F*_1, 9_ = 0.9655, *p* = 0.35), or a day ⨯ condition interaction (*F*_1.574, 14.17_ = 0.1108, *p* = 0.85) for the amount of time spent off nest (Fig. 2B). Control dams spent ∼62% of their time on nest and ∼38% of their time off nest (Fig. 2C) while POE dams spent ∼71% of their time on nest and ∼29% of their time off nest (Fig. 2D). Welch’s unpaired t-tests showed that these percentages were not significant for POE (*M* = 50, *SD* = 29.70) or control (*M* = 50, *SD* = 16.97), *t*_1.590_ = 0.00, *p* > 0.99.

**Figure 2.**
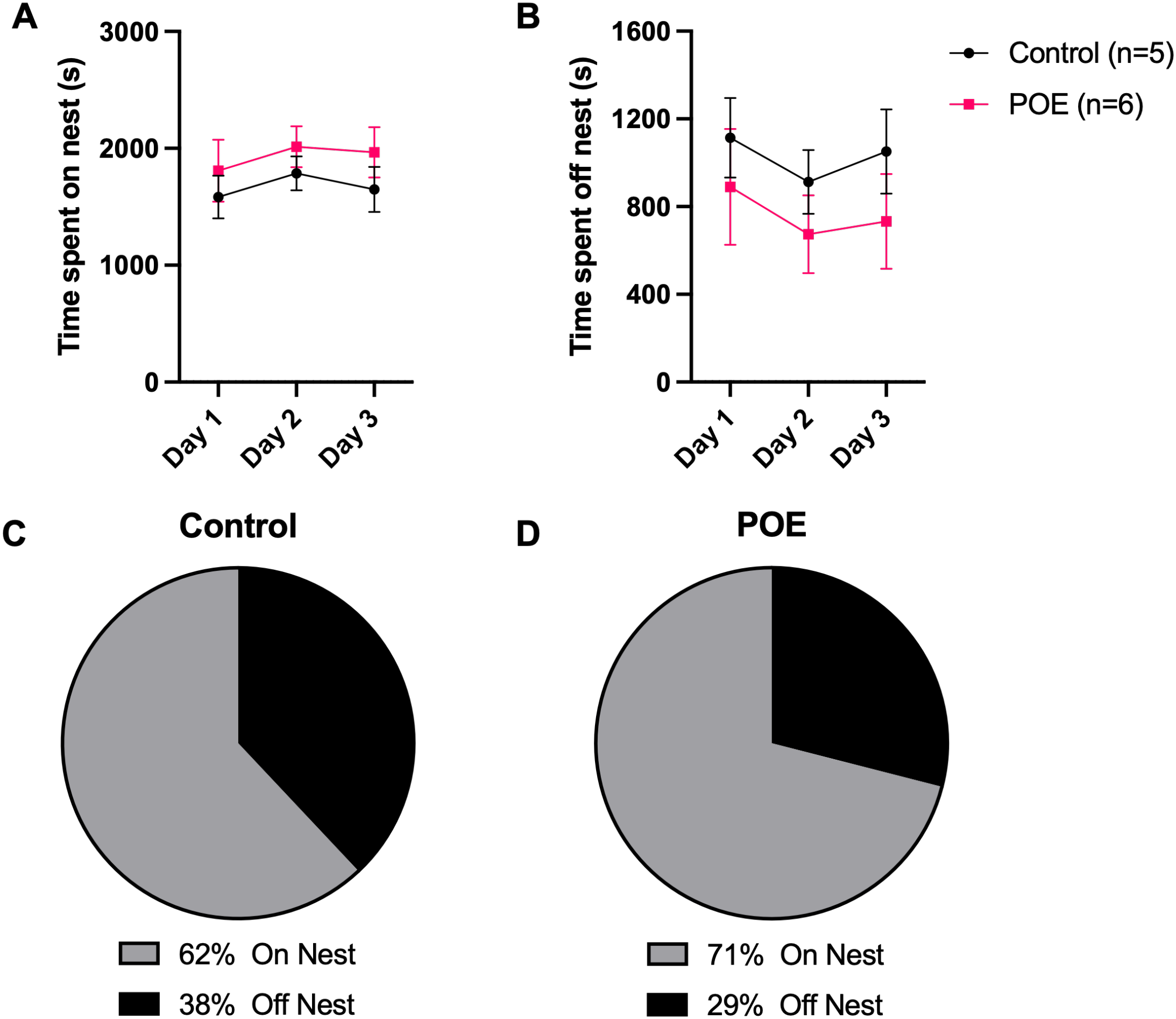
Time spent on or off nest. A) Time spent on nest across days. B) Time spent off nest across days. C) Percentage of time spent on or off nest visualized for control dams. D) Percentage of time spent on or off nest visualized for POE dams. Data are represented as means ± SEMS. POE, Predator Odor Exposure.

### Pup directed behaviors

There was a significant effect of day on time spent licking and grooming pups (*F*_1.803, 16.23_ = 3.808, *p* = 0.048) with Tukey post hoc comparisons revealing an increase between day 1 and day 3 (*p* = 0.037) (Fig. 3A). There was no effect of condition on time spent licking and grooming pups (*F*_1, 9_ = 0.3285, *p* = 0.58). There was also no day condition interaction on time spent licking and grooming pups (*F*_1.803, 16.23_ = 0.1573, *p* = 0.83). There was a significant effect of day on time spent nest building (*F*_1.245, 11.21_ = 7.194, *p* = 0.0169) with Tukey post hoc comparisons revealing a significant decrease between day 1 and day 2 (*p* = 0.0482) as well as day 1 and day 3 (*p* = 0.0242) (Fig. 3B). There was no effect of condition on time spent nest building (*F*_1, 9_= 1.080, *p* = 0.33). There was also no day condition interaction for time spent nest building (*F*_1.245, 11.21_ = 0.649, *p* = 0.47). There were no significant main effects of day (*F*_1.190, 10.71_ = 0.856, *p* = 0.82), condition (*F*_1, 9_ = 0.422, *p* = 0.53), or day condition interaction (*F*_1.190, 10.71_ = 0.017, *p* = 0.93) on time spent sniffing pups (Fig. 3C). There was a significant main effect of day on time spent carrying pups (*F*_1.300, 11.70_ = 5.819, *p* = 0.027), but Tukey post hoc comparisons yielded no significant comparisons (Fig. 3D). There was no effect of condition on time spent carrying pups (*F*_1, 9_ = 0.1023, *p* = 0.76). There was also no day ⨯ condition interaction for time spent carrying pups (*F*_1.300, 11.70_ = 0.2242, *p* = 0.71).

**Figure 3.**
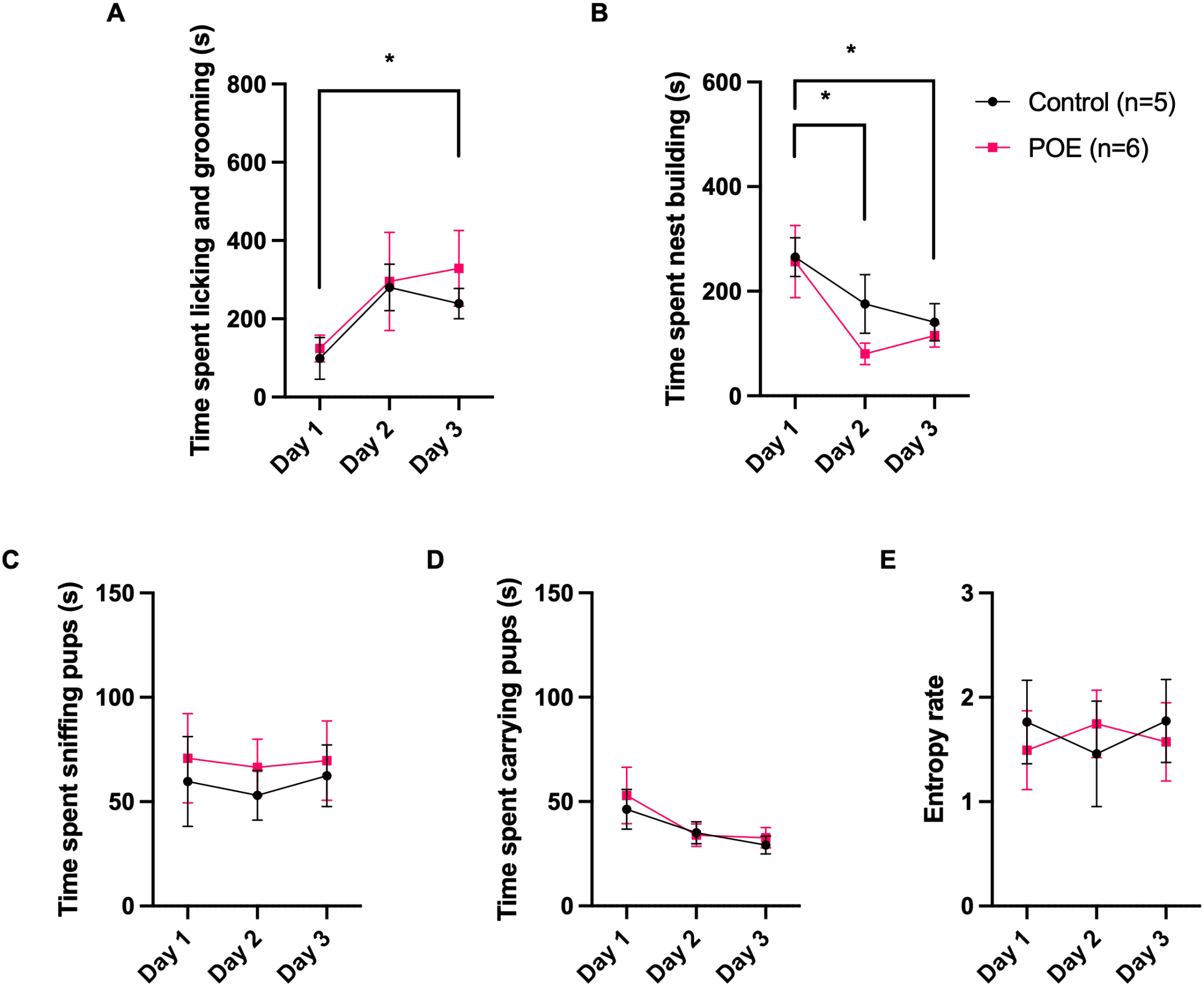
Pup directed behaviors. A) Time spent licking and grooming pups across days. B) Time spent nest building across days. C) Time spent sniffing pups across days. D) Time spent carrying pups across days. E) Behavioral entropy rate across days. *p<.05. Data are represented as means ± SEMS. POE, Predator Odor Exposure.

### Behavioral entropy

There were no significant effects of day (*F*_1.020, 9.177_ = 0.059, *p* = 0.82), condition (*F*_1, 9_ = 0.014, *p* = 0.91), or a day ⨯ condition interaction (*F* _1.020, 9.177_ = 1.052, *p* = 0.33) on behavioral entropy (Fig. 3E).

### Nursing

There were no significant effects of day (*F*_1.664, 14.98_ = 1.572, *p* = 0.67), condition (*F*_1, 9_ = 1.764, *p* = 0.55), or day ⨯ condition interaction (*F*_1.664, 14.98_ = 0.3554, *p* = 0.67) on time spent archback nursing (Fig. 4A). There were no significant effects of day (*F*_1.074, 9.663_ = 3.401, *p* = 0.09), condition (*F*_1, 9_ = 0.3937, *p* = 0.54), or day ⨯ condition interaction (*F*_1.074, 9.663_ = 0.1719, *p* = 0.71) on time spent blanket nursing (Fig. 4B). There ewre also no significant effects of day (*F*_1.000, 9.000_ = 0.8182, *p* = 0.39), condition (*F*_1, 9_ = 0.8182, *p* = 0.39), or day ⨯ condition interaction (*F*_1.000, 9.000_ = 0.8182, *p* = 0.39) on time spent passive nursing (Fig. 4C).

**Figure 4.**
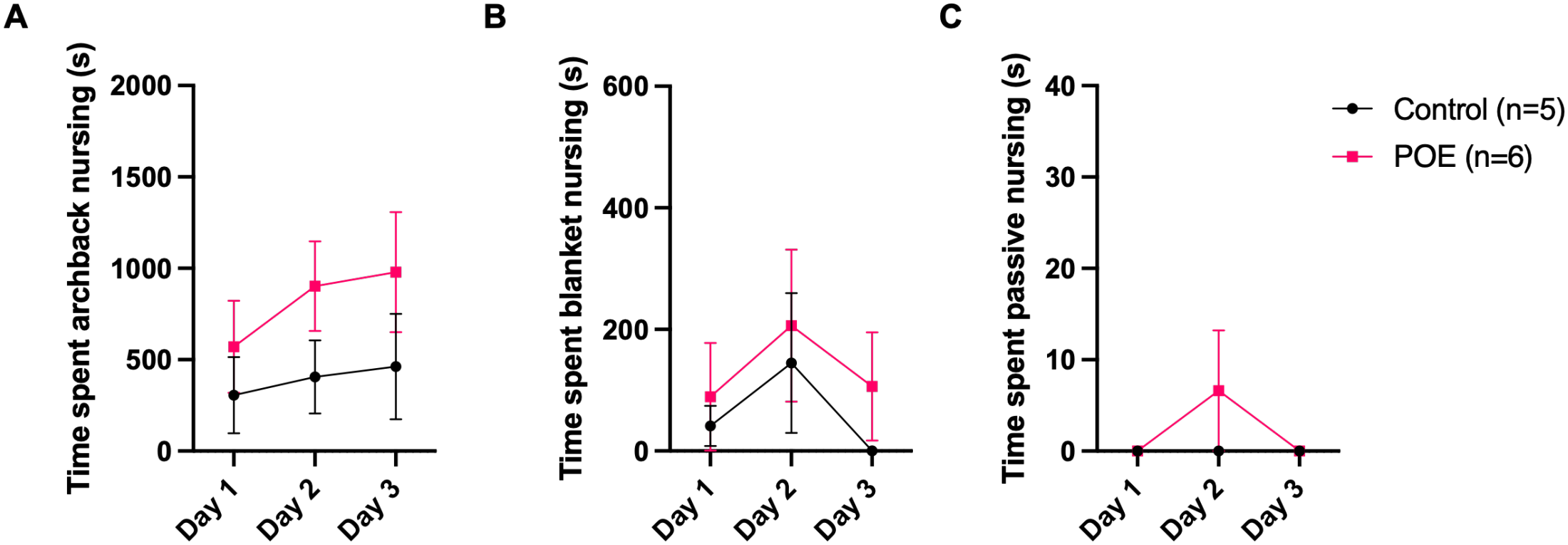
Nursing behavior. A) Time spent archback nursing across days. B) Time spent blanket nursing across days. C) Time spent passive nursing across days. Data are represented as means ± SEMS. POE, Predator Odor Exposure.

### Non-pup directed behaviors

There was a significant effect of day on instances of rearing (*F*_1.074, 9.664_ = 8.932, *p* = 0.0131) with Tukey post hoc comparisons revealing a significant decrease in instances between day 1 and day 2 (*p* = 0.0491) and day 1 and day 3 (*p* = 0.0073) (Fig. 5A). There were no significant effects of condition on rearing (*F*_1, 9_ = 0.04185, *p* = 0.84), nor day ⨯ condition interactions for rearing (*F*_1.074, 9.664_ = 0.1723, *p* = 0.70). There were no significant effects of day (*F*_1.476, 13.29_ = 1.912, *p* = 0.19), condition (*F*_1, 9_ = 0.243, *p* = 0.63), or day ⨯ condition interactions (*F* _1.476, 13.29_ = 2.491, *p* = 0.13) on time spent self-grooming (Fig. 5B).

**Figure 5.**
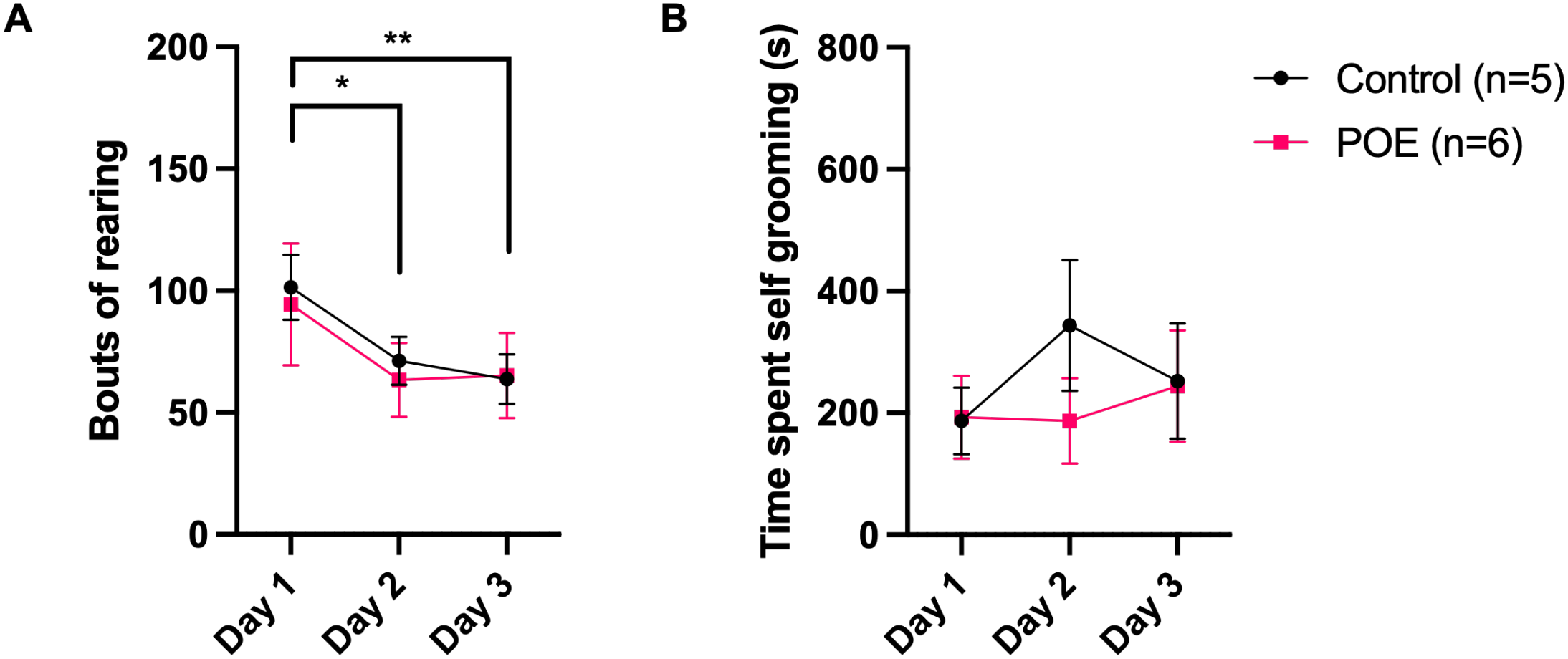
Non-pup directed behavior. A) Number of bouts of rearing across days. B) Time spent self-grooming across days. *p<.05, **p<.01. Data are represented as means ± SEMS. POE, Predator Odor Exposure.

## Discussion

Neonatal POE produces lasting decreases in social reward motivation (Cuarenta et al., 2021; Stricklin et al., 2026) and impacts the developing brain (Chang et al., 2019; Cuarenta et al., 2022; Kigar et al., 2017), but it was not yet known if POE altered maternal care, potentially contributing to the observed impacts. The present study examined if neonatal POE altered subsequent maternal care behaviors. Overall, our results demonstrate that dams do not alter their maternal care in response to pups receiving POE. There were no significant effects of POE on latency to retrieve, time spent on or off nest, sniffing, time spent carrying pups during the 45 minutes of maternal care, nursing, or behavioral entropy. We predicted that POE would result in increased pup-directed behaviors, but the only significant effect of condition was seen on time spent carrying pups during retrieval on day 1, where control dams spent significantly more time carrying pups during retrieval than POE dams. There were also significant effects of day on time spent carrying pups to retrieval, time spent nest building, time spent licking and grooming, and rearing. This suggests that the impacts we observe from our neonatal POE model are due to the exposure itself on the pups, and not any alterations in maternal care.

We measured latency to retrieve as the time it took for the dam to return all pups to the nest. Timely retrieval is crucial for pup warmth, safety, and access to maternal care, as delays in retrieval could alter the care pups receive while outside the nest. Latency to retrieve all pups to the nest did not differ between conditions and remained similar across days. However, control dams spent more time carrying pups during retrieval on day 1 than days 2 and 3, potentially indicating increased retrieval efficiency with repeated separation. In contrast, POE dams showed consistent carrying duration during retrieval of pups across all days. One possibility is that the decrease in carrying duration across days in control dams may reflect differences in pup vocalizations. Previous research has shown that maternal separation can alter the quantity, acoustic properties, and duration of ultrasonic vocalizations from rat pups (Yuksel et al., 2026), and limited bedding and nesting can increase both ultrasonic and audible vocalizations from mice pups (Heun-Johnson & Levitt, 2016). Future studies examining pup vocalizations following POE may help clarify these differences on day 1 in maternal retrieval.

There were no differences in nest building, licking and grooming, and rearing in POE vs. Control dams. However, these measures changed for both groups across the 3 days. We observed that time spent nest building decreased across days, indicating that dams may become more efficient at this behavior as the days progress. Time spent licking and grooming increased between day 1 and day 3, which may be reflective of the dams having more time to devote to this behavior because some other behaviors, like nest building and rearing, are decreasing. There was also a significant effect of day on rearing such that both POE and control dams reared more on day 1 than day 2 or day 3. This can largely be attributed to the novelty of the plexiglass cage lid that was used for behavior recording. To enable recording of the cage, the normal food hopper and lid were removed and replaced for the duration of the recording with a clear plexiglass testing lid. On day 1 especially, dams from both conditions spent a lot of time rearing and sniffing or chewing on the lid. Future experiments should allow the dams additional time to acclimate to the novel testing lid.

Previous research has found that at PND33, POE females have significantly higher body weights than control females (Kigar et al., 2024). We did not find any effects of POE of time spent nursing, including archback, passive, and blanket nursing subtypes. This suggests that differences in adolescent body weight are unlikely to be explained by changes in the amount of time dams spend nursing pups. It’s possible POE increases adolescent female body weight by altering other mechanisms, like the HPA axis, which is known to be associated with weight gain (Lengton et al., 2025). We hypothesized that dams of POE litters would engage in more pup sniffing in response to residual odor following POE. If dams were able to detect the odor, we would expect increased sniffing as they investigated the unfamiliar odor. Alternatively, dams may find the predator odor aversive and spend less time sniffing their pups. However, we found no differences in time spent sniffing pups between POE and control dams, suggesting that dams likely do not detect residual predator odor on their pups. This finding is particularly relevant to our model, as POE is designed to expose pups to predation threat without directly exposing dams. The absence of changes in pup sniffing provides further support that dams are not responding to the predator odor itself. We also found no differences in behavioral entropy, which is characteristic of other ELA models that alter maternal care such as LBN that which produces unpredictable, fragmented care (Baram et al., 2012; Goodwill et al., 2019; Ivy et al., 2008). One limitation of this study is that we only observed maternal care for 45 minutes following POE on each of the three exposure days, totaling 135 minutes of observation time for each litter. Thus, our observations do not capture maternal behavior throughout the neonatal period, and changes in maternal behavior may emerge at a later postnatal date.

Different models capture distinct dimensions of ELA, which is an important consideration when evaluating these results. Previous research on ELA often utilizes limited bedding and nesting or maternal separation models (Eck et al., 2020; Ivy et al., 2008; Molet et al., 2014), which are conceptualized as resource scarcity and neglect. Neonatal POE captures a specific dimension of ELA, the stress of perceived predation threat. Threat predation is a stressor for many species, highlighting the ethological relevance of our POE model. Encounters with a predatory threat activate stress-response systems and prompt defensive behavior in response in a number of mammalian prey species including rats, mice, voles, rabbits, deer, and sheep (Apfelbach et al., 2005; Blanchard et al., 2003; Hegab & Wei, 2014), even if this encounter does not actually pose a threat, like our POE model. The ability to distinguish threatening from non-threating stimuli is particularly relevant to post-traumatic stress disorder, which is characterized in part by altered fear responses to both types of stimuli (Bakshi et al., 2000). Our neonatal model of POE is applicable to other species of prey animals, as the exact odors used can be representative of the natural predators of that species.

Previous research also demonstrates that most models of ELA change some aspect of maternal care (Baram et al., 2012; Brunson et al., 2005; Eck et al., 2020; Goodwill et al., 2019; Ivy et al., 2008; Liu et al., 1997; Molet et al., 2014), but the present study provides evidence that POE does not alter maternal care behavior in a meaningful way. This can largely be attributed to most ELA models impacting the environment that is shared by both the dam and the pups, meaning that the developmental impacts of these models are due to a combination of direct effects on the pups and indirect effects resulting from alterations to maternal care. Neonatal POE is also a unique model in that exposure is restricted to the pups only, and not the dam. This model provides an opportunity to investigate the effects of neonatal threat on development independent of alterations in maternal care.

## Author contributions

**Victoria R. Riesgo:** Investigation, Formal analysis, Visualization, Writing – original draft. **Margie C. Stricklin:** Investigation, Writing - original draft. **Hannah E. Carney:** Investigation. **Judy E. Nyakoa:** Investigation. **Cecelia J. Crandall:** Investigation. **Debra A. Bangasser:** Funding acquisition, Supervision, Writing - review & editing. **Amelia Cuarenta:** Conceptualization, Methodology, Investigation, Funding acquisition, Project administration, Writing - review & editing.

## Acknowledgements

This work was supported by the National Institutes of Health (R21 DA059966, R01 DA049837, R01 DA056534, R34 DA061483, R21 DA062844, K99/R00 DA060266), the National Science Foundation (IOS-2313253, IO-2208822), and the Wilczynski-Georgia Research Alliance Future Faculty Fellowship in Neuroscience Award. The content is solely the responsibility of the authors and does not necessarily represent the official views of the NIH or NSF. A special thank you is extended to Erin Harris for 3D printing the chambers and platforms used for POE, and Dario Aspesi for providing the R code for behavioral entropy rate.

